# Three-dimensional label-free histological imaging of whole organs by microtomy-assisted autofluorescence tomography

**DOI:** 10.1101/2021.09.07.459253

**Authors:** Yan Zhang, Lei Kang, Wentao Yu, Victor Tsz Chun Tsang, Terence T. W. Wong

## Abstract

Three-dimensional (3D) histology is vitally important to characterize disease-induced tissue heterogeneity at the individual cell level. However, it remains a scientific challenge for both high-quality 3D imaging and volumetric reconstruction. Here we propose a label-free, automated, and ready-to-use 3D histological imaging technique, termed microtomy-assisted autofluorescence tomography with ultraviolet excitation (MATE). With the combination of block-face imaging and serial microtome sectioning, MATE can achieve rapid and label-free imaging of paraffin-embedded whole organs at an acquisition speed of 1 cm^3^ per 4 hours with a voxel resolution of 1.2 × 1.2 × 10 μm^3^. We demonstrate that MATE enables simultaneous visualization of cell nuclei, fiber tracts, and blood vessels in mouse/human brains without tissue staining or clearing. Moreover, diagnostic features, such as nuclear size and packing density, can be quantitatively extracted with high accuracy. MATE is augmented to the current slide-based 2D histology, holding great promise for facilitating histopathological interpretation at the cell level to analyze complex tissue heterogeneity in 3D.

**Significance Statement:** Conventional 3D histology based on spatial registration of serial histochemically-stained thin tissue slices is fundamentally labor-intensive and inaccurate. Here, we propose a rapid and label-free 3D histological imaging technique (i.e., MATE) that enables high-resolution imaging of complex whole organs without tissue staining or clearing. MATE is fully automated to provide a series of distortion- and registration-free images with intrinsic absorption-based contrast, demonstrating great potential as a routine tissue analysis tool that can seamlessly fit into the current clinical practice to facilitate the applications of histopathological interpretation at the subcellular level.

## Introduction

Conventional slide-based two-dimensional (2D) histology is commonly used clinically. However, it fails to provide sufficient representation of large and bulky tissues. On the contrary, 3D histology is vital to study structural changes volumetrically at the cell level, allowing accurate diagnosis of continuously spreading tumor strands (1). Yet, conventional 3D histology (2, 3) based on image registration of serial histochemically-stained thin tissue slices is fundamentally slow, labor-intensive, and inaccurate due to the inevitable tissue ruptures during slide preparation, which further poses a challenge for the registration of adjacent slices during volumetric reconstruction (4). Non-invasive tomographic imaging techniques, including optical projection tomography (5), microscopic magnetic resonance imaging (6), and X-ray micro-computed tomography (μCT) (7), enable rapid 3D imaging of tissue’s anatomical structures with resolution ranging from sub-micrometer to tens of micrometers. However, they are not suitable for probing molecular targets as desired in clinical standard histopathology. Furthermore, it is not possible to achieve high-quality imaging of soft tissues in μCT without X-ray-absorbing staining agents (8).

Modern advancements in optical microscopy, data processing, and 3D visualization pave a way for multi-scale 3D analysis of complex tissue networks at microscopic resolution. However, due to light scattering and absorption, traditional optical imaging techniques (9–11) can only image tens to hundreds of microns deep into the tissue, hindering their applications of 3D imaging of whole organs. To increase the accessible imaging depth, light-sheet fluorescence microscopy (LSFM) of chemically cleared tissues (12–15) has been extensively utilized for rapid volumetric imaging. However, the imaging depth is inherently traded with the lateral resolution in LSFM systems, and the imaging quality significantly degrades with depth due to inhomogeneous tissue clearing (16). In addition, slow diffusion and limited penetration of fluorescent probes into the cleared tissues is a common issue (17). Alternatively, imaging systems based on the integration of serial mechanical sectioning provide a new strategy to overcome the depth limitation. Serial two-photon tomography (STPT) (18, 19), block-face serial microscopy tomography (FAST) (20), wide-field large-volume tomography (WVT) (21), micro-optical sectioning tomography (MOST) (22), and fluorescence MOST (fMOST) (23) are all examples of this approach. With the combination of ultra-thin sectioning and line-scan imaging on a knife edge, MOST and fMOST enable to image resin-embedded whole mouse brains at 0.33 × 0.33 × 1 μm^3^ voxel resolution within ~10 days (22). In contrast, block-face imaging systems (i.e., STPT, FAST, and WVT) are implemented by alternating cycles of en-face imaging of a tissue block and mechanical removal of the imaged surface. These systems considerately accelerate the 3D imaging pipeline by adopting a relatively large slicing interval, and the captured serial block-face images are inherently registered, ensuring an accurate volumetric reconstruction. For instance, by using spinning disk confocal microscopy, FAST demonstrates an unprecedented speed that can achieve whole-brain imaging at 0.7 × 0.7 × 5 μm^3^ voxel resolution within 2.4 hours (20). The aforementioned systems are powerful tools to study 3D microanatomical structures of large tissues. However, they all require well-regulated and time-consuming protocols regarding tissue preparation, such as clearing (24), staining, and embedding (25) of an intact organ, making it challenging to be integrated into the current histology workflow. Although microtomy-assisted photoacoustic microscopy (26) enables label-free whole-brain imaging at 0.91 × 0.91 × 20 μm^3^ voxel resolution, the long acquisition time (~15 days with a 200-μm slicing interval) eventually hinders its utility as a routine tissue analysis approach.

To this end, we propose a label-free, automated, and ready-to-use 3D histological imaging method, termed microtomy-assisted autofluorescence tomography with ultraviolet (UV) excitation (MATE). Rich endogenous fluorophores (27), including reduced nicotinamide adenine dinucleotide, flavin coenzymes, structural proteins, aromatic amino acids, porphyrins, and lipopigments naturally fluoresce with deep-UV excitation (28), generating an intrinsic absorption-based contrast for label-free histological imaging. In addition, moderate optical sectioning strength (~10 μm (29)) provided by UV surface excitation (30–32) allows MATE to be integrated with block-face imaging and microtome sectioning to enhance the accessible imaging depth. The automated MATE system provides serial registration-free block-face images of formalin-fixed and paraffin-embedded (FFPE) whole organs without tissue staining or clearing, achieving rapid 3D imaging of the whole mouse brain at 1.2 × 1.2 × 10 μm^3^ voxel resolution within 3.5 hours, generating a brain atlas like dataset which consists of 900 serial coronal sections with an overall data size of ~300 gigabytes. Here, we demonstrate that MATE enables simultaneous visualization of various anatomical structures, including cell nuclei, fiber tracts, and blood vessels in mouse/human brains with high fidelity. Moreover, diagnostic features, such as nuclear size and packing density, can be quantitatively extracted from MATE images with high accuracy. The high speed, low cost, and ease-of-use features of MATE demonstrate its great potential as a routine tissue analysis tool that can seamlessly fit into the current histology workflow, facilitating the applications of histopathological interpretation at the cell level to analyze complex tissue networks.

## Results

### MATE system for whole-organ imaging and sectioning

MATE is configured in reflection mode to accommodate tissues with any size and thickness (Fig. 1). A UV light-emitting diode (LED) with 285-nm wavelength is implemented for the excitation of endogenous fluorophores based on intrinsic absorption. With oblique illumination, we circumvent the use of UV-transmitting optics and fluorescence filters because the backscattered UV light is naturally blocked by the glass objective and tube lens which are spectrally opaque at 285 nm. The FFPE block specimen is clamped towards the objective lens by an adjustable sample holder, and rigidly mounted on a three-axis motorized stage to control both the scanning for in-plane mosaic imaging and tissue sectioning by a lab-built microtome (inset of Fig. 1). The alternating cycles of en-face imaging of a tissue block and mechanical removal of the imaged surface are repeated until the whole block specimen is captured. With the synchronization of block-face imaging and microtome sectioning, the automated MATE imaging system provides serial distortion- and registration-free z-stack images evenly spaced at a 10-μm interval, enabling rapid and label-free 3D imaging at an acquisition speed of 1 cm^3^ per 4 hours with a measured lateral resolution of 1.2 μm (Supplementary Fig. 1).

**Figure 1.**
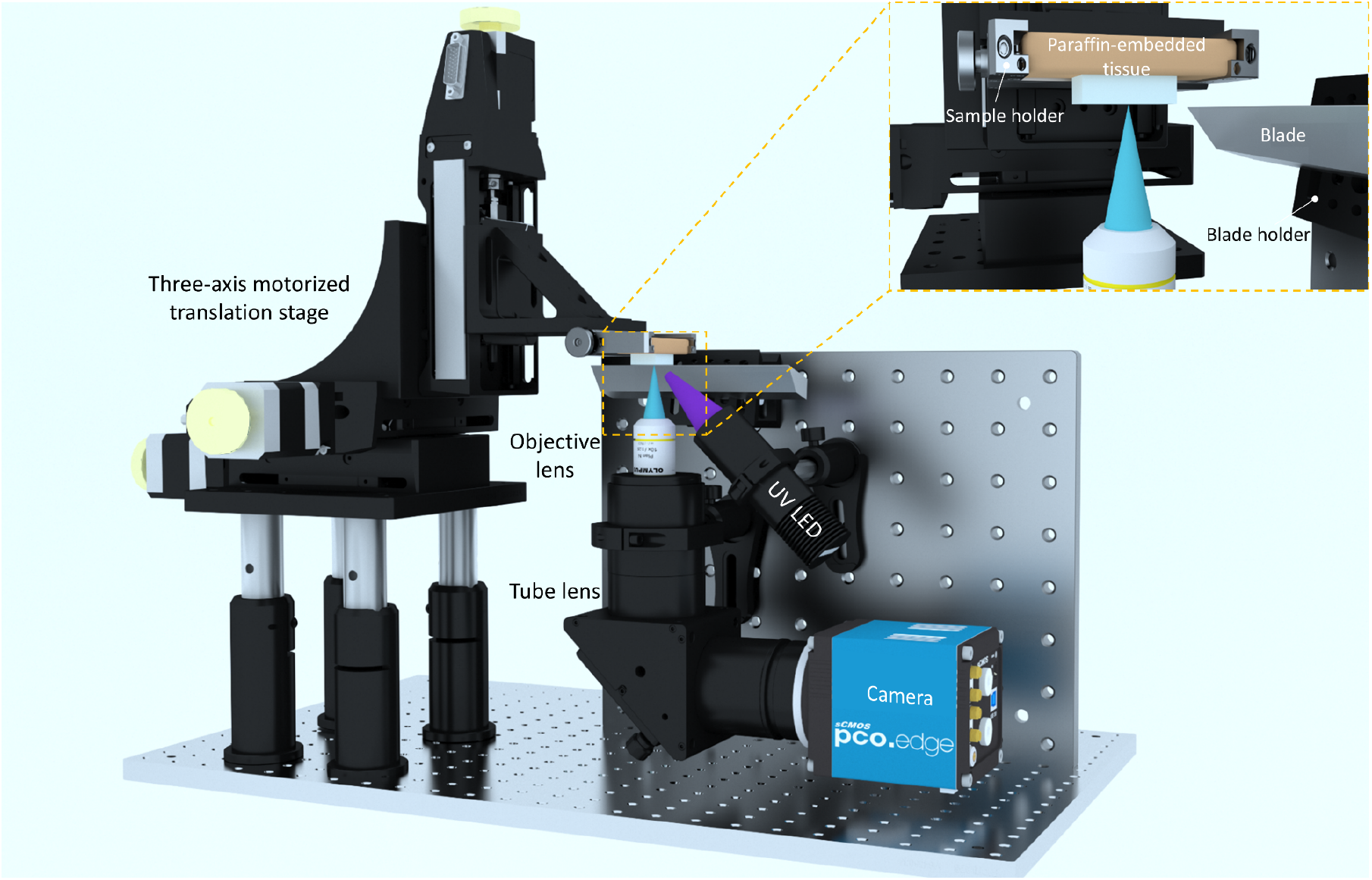
Schematic of the MATE system for whole-organ imaging and sectioning. The UV light from a UV LED is focused onto the bottom surface of a block specimen, which is clamped towards the objective lens by an adjustable sample holder, and rigidly mounted on a three-axis motorized translation stage to control both the scanning for in-plane imaging and tissue sectioning. The excited intrinsic fluorescence signals from the exposed tissue surface are collected by an objective lens, refocused by a tube lens, and subsequently detected by a monochromatic camera. The imaged surface is then sectioned by a lab-built microtome (blade) so that a new surface is exposed for imaging. This process is repeated until the entire tissue block is imaged.

### MATE imaging of an intact FFPE mouse brain block

The performance of MATE is initially validated by imaging FFPE thin slices of mouse kidney/liver tissues (Supplementary Fig. 2). The microtome-sectioned thin tissue slices are deparaffinized to minimize paraffin-induced fluorescence for MATE imaging. A variety of anatomical structures, including renal tubules, Bowman’s space, and glomerulus in mouse kidney (Supplementary Fig. 2b–d), and hepatocytes, hepatic sinusoids, and erythrocytes in mouse liver (Supplementary Fig. 2i–k) are clearly identified. After MATE imaging, the slices are histochemically stained by hematoxylin and eosin (H&E), and imaged with a bright-field microscope to obtain the corresponding histological images for comparison (Supplementary Fig. 2e–g, l–n). Multiple similarities are revealed in MATE and H&E-stained images, despite that the nucleolar structures in mouse liver are less visible in MATE images. Pearson correlation coefficient of 0.9 is calculated from Supplementary Fig. 2b and e, validating the feasibility of using tissue’s autofluorescence as an intrinsic contrast for label-free histological imaging.

The moderate optical sectioning strength provided by UV surface excitation enables MATE to be utilized for slide-free histological imaging, which is validated by imaging an FFPE mouse brain block (Supplementary Fig. 3 and Supplementary Video 1). The mouse brain block is imaged by the MATE system (Supplementary Fig. 3a–f), and subsequently sectioned at the surface by a microtome with 7-μm thickness to obtain the corresponding H&E-stained images for comparison (Supplementary Fig. 3g–l). Although the slide-based histological images cannot exactly replicate the imaged surface by MATE due to the different imaging depth and tissue distortion, the histological features, including isocortex, hippocampus, thalamus, and cortical amygdala are still remarkably similar. Nuclear features, such as cross-sectional area and intercellular distance, are extracted from MATE and H&E-stained images (e.g., Supplementary Fig. 3b and h) for quantitative analysis. The difference between these two distributions of nuclear features is evaluated by Wilcoxon rank-sum test under a significance level of 0.05. Our results suggest that the nuclear features extracted from MATE agree fairly well with the H&E-stained histological image (Supplementary Fig. 3m,n), validating the accuracy of the information that can be provided by MATE.

To determine the optimal sectioning interval for 3D whole-brain imaging, the imaging depth of MATE is estimated by leveraging the fact that nuclear count is linearly proportional to the accessible tissue imaging depth. First, an FFPE mouse brain block is imaged by MATE (Supplementary Fig. 4a). Then, the same tissue block is consecutively sectioned at the block surface with 7-μm thickness, and subsequently stained with H&E to obtain 3 adjacent slices (Supplementary Fig. 4b–d), which reveal the nuclear distribution within a depth range of 21 μm in total. The ratio of the nuclear count in the H&E-stained images within a given depth range to that in the MATE image is calculated to be closest to unity for a depth range of 7 μm (Supplementary Fig. 4i), suggesting that the imaging depth of MATE is within 10 μm.

The full capacity of MATE for 3D high-resolution imaging is demonstrated in Fig. 2. An intact FFPE mouse brain block (top inset of Fig. 2a) is imaged by MATE at 10-μm sectioning interval within 3.5 hours, generating a brain atlas-like dataset consisting of 900 serial registration-free coronal sections with an overall size of ~300 gigabytes (Supplementary Video 2). The 3D model with orthogonal views of the imaged whole brain is shown in Fig. 2a. Anatomical structures including cerebellum, cerebral cortex, hippocampus, thalamus, and olfactory bulb are clearly visualized in the coronal (Fig. 2b), transverse (Fig. 2c), and sagittal (Fig. 2d) images. The densely packed cell nuclei in the hippocampus (Fig. 2e–g) can be resolved individually with a lateral resolution of 1.2 μm. Each nucleus in the coronal section can be localized by Jerman’s spherical enhancement filter (33) which is based on the ratio of multiscale Hessian eigenvalues, generating a nuclear density map across the whole coronal section (Fig. 2i and Supplementary Fig. 5). Note that the nuclear count is not absolutely accurate because the vascular cross sections, which present similar brightness to cell nuclei in the MATE images, are also extracted by Jerman’s filter. We believe that integrating with advanced segmentation neural networks would enable a more faithful statistical analysis. Ten representative coronal sections with the relative positions marked in a transverse view (Fig. 2j) are shown in Fig. 2k–t. Volumetric rendering of the hippocampus and cerebellum indicated by yellow solid and blue dashed regions in Fig. 2j are demonstrated in Fig. 2u and v, respectively.

**Figure 2.**
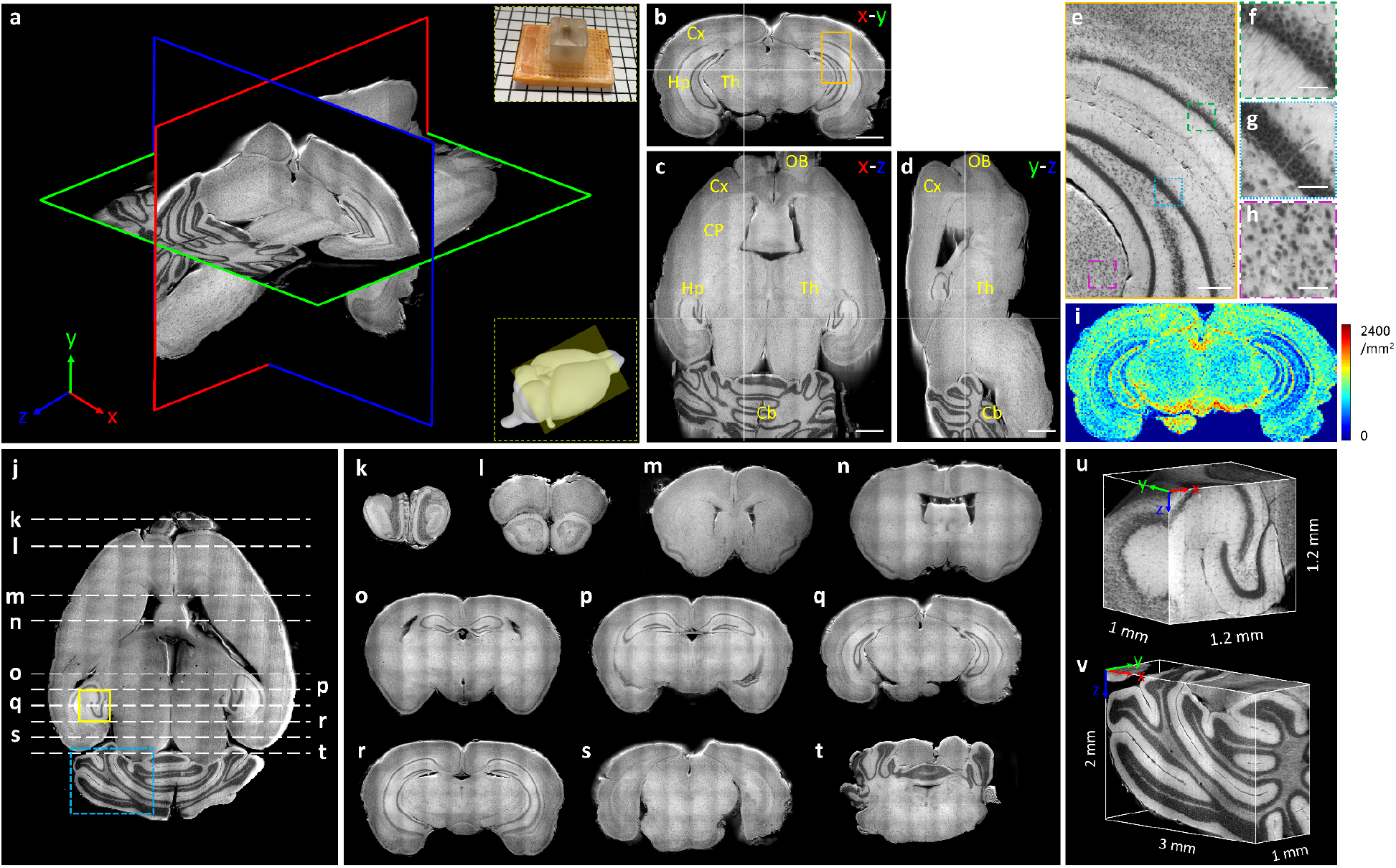
3D label-free MATE imaging of an intact mouse brain embedded in an FFPE block. **a**, A 3D model with orthogonal views of the imaged whole mouse brain. The top inset shows the photograph of the specimen, and the bottom inset indicates the imaging area of a 3D mouse brain model (yellow shaded region). **b**–**d**, The corresponding coronal view (*x-y* plane), transverse view (*x-z* plane), and sagittal view (*y-z* plane) of the imaged brain, respectively. Crosshairs are linked in three orthogonal views, and anatomical structures are annotated in each view. OB, olfactory bulb; Cx, cerebral cortex; CP, caudate putamen; Hp, hippocampus; Th, thalamus; Cb, cerebellum. **e**, A zoomed-in MATE image of the orange solid region in b. **f**–**h**, Zoomed-in MATE images of green, blue, and magenta dashed regions in e, respectively. **i**, A nuclear density map calculated from b. **j**, A transverse view of the brain volume. **k**–**t**, Ten representative coronal sections with the relative positions marked in j. **u,v**, Volumetric rendering of yellow solid and blue dashed regions in j, respectively. Scale bars, 1 mm (b–d), 200 μm (e), and 50 μm (f–h).

Figure 3 shows a variety of histological features extracted collectively from the imaged mouse brain block, showing in the coronal view. Fiber tracts (Fig. 3a–c), blood vessels (Fig. 3d,e), ventricular system (Fig. 3f,g), and cell populations (Fig. 3h–o) located at different regions across the whole brain are simultaneously visualized in MATE images with high fidelity in a label-free manner. This unique capability of MATE enables comprehensive 3D histopathological analysis of complex whole organs, thus holding great promise for the investigation of structural connectivity and heterogeneity involved in many disease processes. To further explore the effect of different embedding materials, a formalin-fixed and agarose-embedded mouse brain is sectioned by a vibratome with 200-μm thickness for MATE imaging. Two representative coronal sections are shown in Supplementary Fig. 6a–e, and the corresponding features extracted from a paraffin-embedded mouse brain are utilized for comparison (Supplementary Fig. 6f–h). An obvious distinction is observed that autofluorescence from fiber tracts (e.g., internal capsule and cerebral peduncle) and myelin-rich globus pallidus is significantly quenched by the tissue-infiltrative paraffin, resulting in a homogenous intensity distribution across the whole brain. In addition, tissue shrinkage is inevitable during paraffin embedding, with a reported percentage of shrinkage ranging from 10% to 27% (34, 35). Although agarose-embedded tissues may be more suitable for MATE imaging, the slicing quality is far inferior to that with paraffin-embedded tissues at a 10-μm interval. We believe that integrating a vibrating blade (i.e., vibratome) with an agarose-embedded tissue can further improve the slicing and imaging quality.

**Figure 3.**
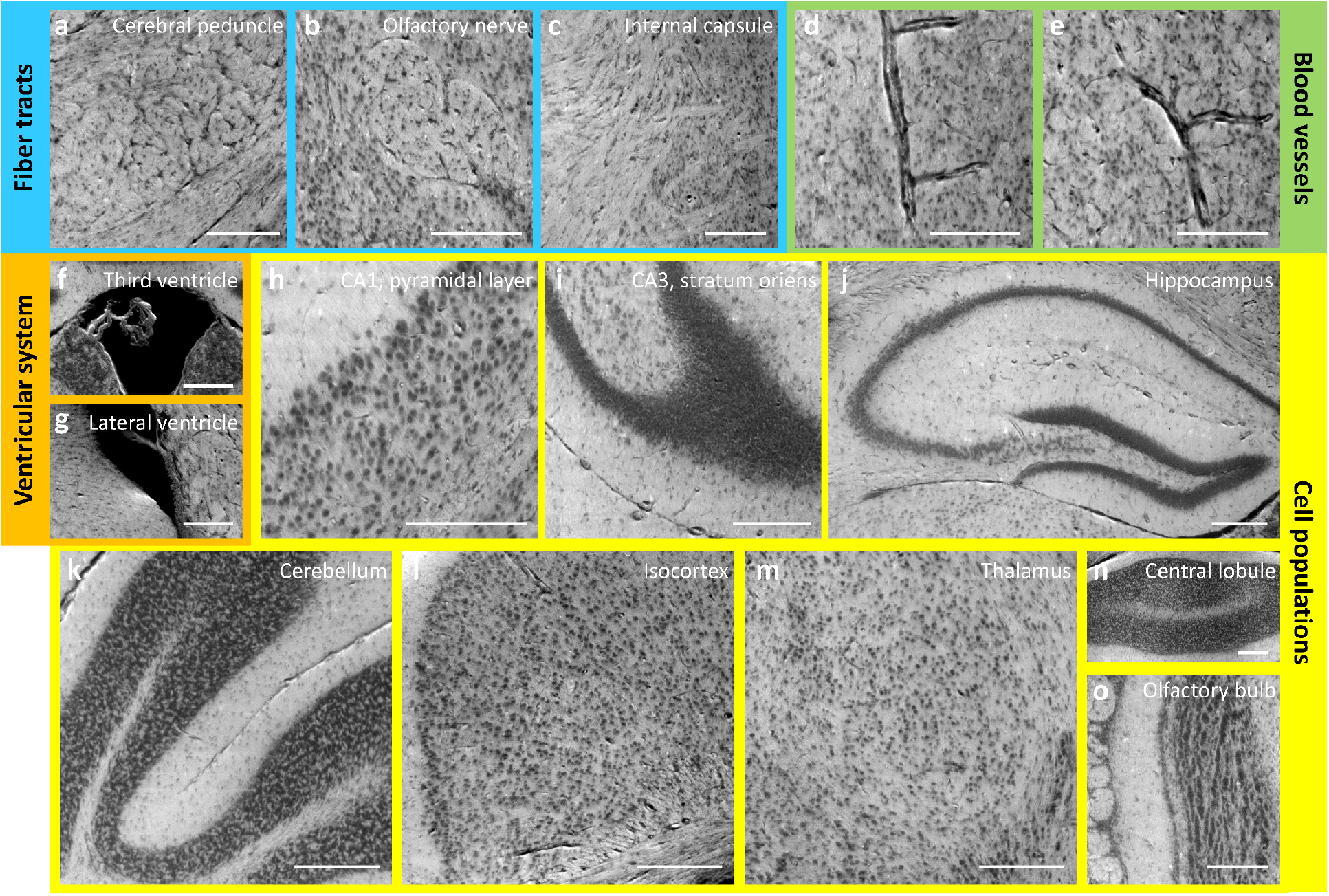
Image gallery of features extracted from label-free MATE images of the FFPE mouse brain block. **a**–**c**, Fiber tracts. **d,e**, Blood vessels. **f,g**, Ventricular system. **h**–**o**, Cell populations located at different regions across the whole brain. All features are shown in a coronal view. Scale bars, 200 μm.

### MATE imaging of an FFPE human brain block

High-resolution 3D MATE imaging of an FFPE human brain block is shown in Fig. 4. The block specimen with a dimension of 11 × 4.5 × 1.5 mm^3^ is imaged by MATE at a 10-μm sectioning interval within 30 minutes, generating a dataset consisting of 150 serial sections with an overall size of ~35 gigabytes (Supplementary Video 3). The 3D model of the imaged volume is illustrated in Fig. 4a, with its corresponding orthogonal views shown in Fig. 4b–d. Volumetric rendering of a portion of the cerebellar cortex (Fig. 4e) outlines a clear boundary between the granular layer (GL) and molecular layer (ML). With cellular resolution, the densely packed granule cells (Fig. 4f) are resolved individually. Structural connectivity in the white matter is clearly revealed in Fig. 4h, where fiber pathways with the intertwined cell populations are visualized with high resolution (Fig. 4i,j and Supplementary Video 4). 3D projection of the region marked in Fig. 4b is color-coded in depth to characterize the vascular network of the human brain in space (Fig. 4k).

**Figure 4.**
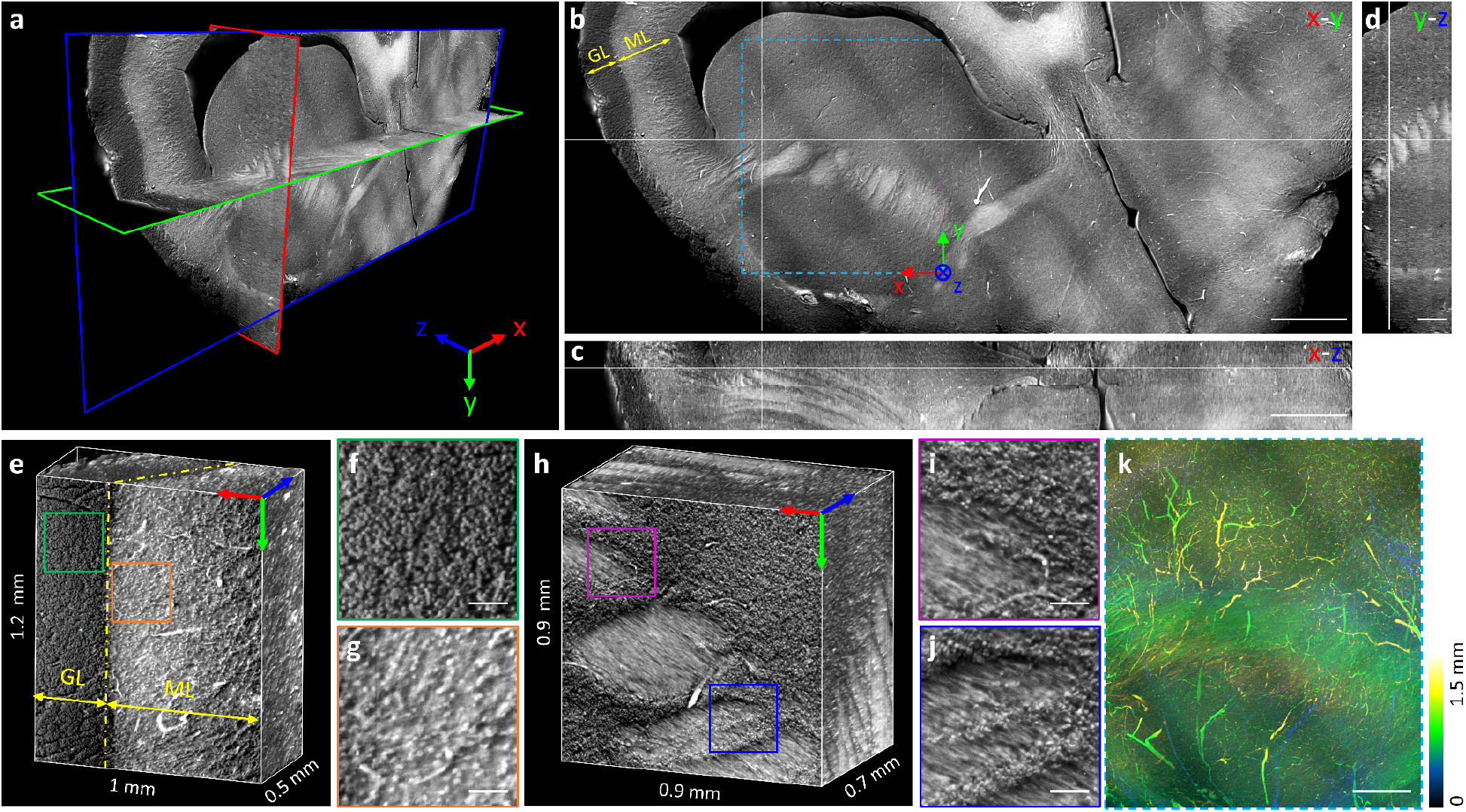
3D label-free MATE imaging of an FFPE human brain block. **a–d**, A 3D model generated by MATE imaging with its corresponding orthogonal views of an imaged human brain block. Crosshairs are linked in three orthogonal views. **e**, Volumetric rendering of a portion of the cerebellar cortex, with the boundary between granular layer (GL) and molecular layer (ML) marked by a yellow dashed line. **f,g**, Zoomed-in MATE images of green and orange solid regions in e, respectively. **h**, Volumetric rendering of fiber pathways in the white matter. **i,j**, Zoomed-in MATE images of magenta and blue solid regions in h, respectively. **k**, Vascular network with color-coded depth obtained by 3D projection of the blue dashed region in b. Scale bars, 1 mm (b,c), 50 μm (f,g,i,j), and 500 μm (d,k).

Figure 5 shows a collection of histological features extracted from the human brain dataset (Fig. 5a–i). Several thin tissue slices were consecutively sectioned from the block surface, and subsequently stained to obtain the closest H&E-stained histological images, which can serve as references of the imaged brain volume. The corresponding features extracted from these H&E-stained images are shown for comparison (Fig. 5j–r). Although these features cannot be identical to that in MATE images due to the tissue perturbation during slide preparation, the cellular distributions are still remarkably similar. For instance, the boundary between the granular layer and molecular layer in the cerebellar cortex is clearly distinguished in both MATE and H&E-stained images (Fig. 5a,b,j,k). The densely packed granule cells and erythrocytes are resolved with high correspondence (Fig. 5c,d,l,m). Other anatomical structures, including choroid plexus (Fig. 5e), neuropil (Fig. 5f), blood vessels (Fig. 5g), and myelinated fibers (Fig. 5h,i) are simultaneously visualized by MATE in a label-free setting that otherwise would require tissue clearing and multiple fluorescent probes for revealing them as mentioned in the previously reported method (36). MATE enables label-free histological imaging of complex tissue networks by non-specific excitation of various biomolecules with minimal tissue preparation, demonstrating its unique superiority as a routine 3D imaging platform. MATE can seamlessly fit into the current clinical practice to facilitate comprehensive histopathological interpretation at the cell level, hence understanding organs in 3D instead of 2D slices.

**Figure 5.**
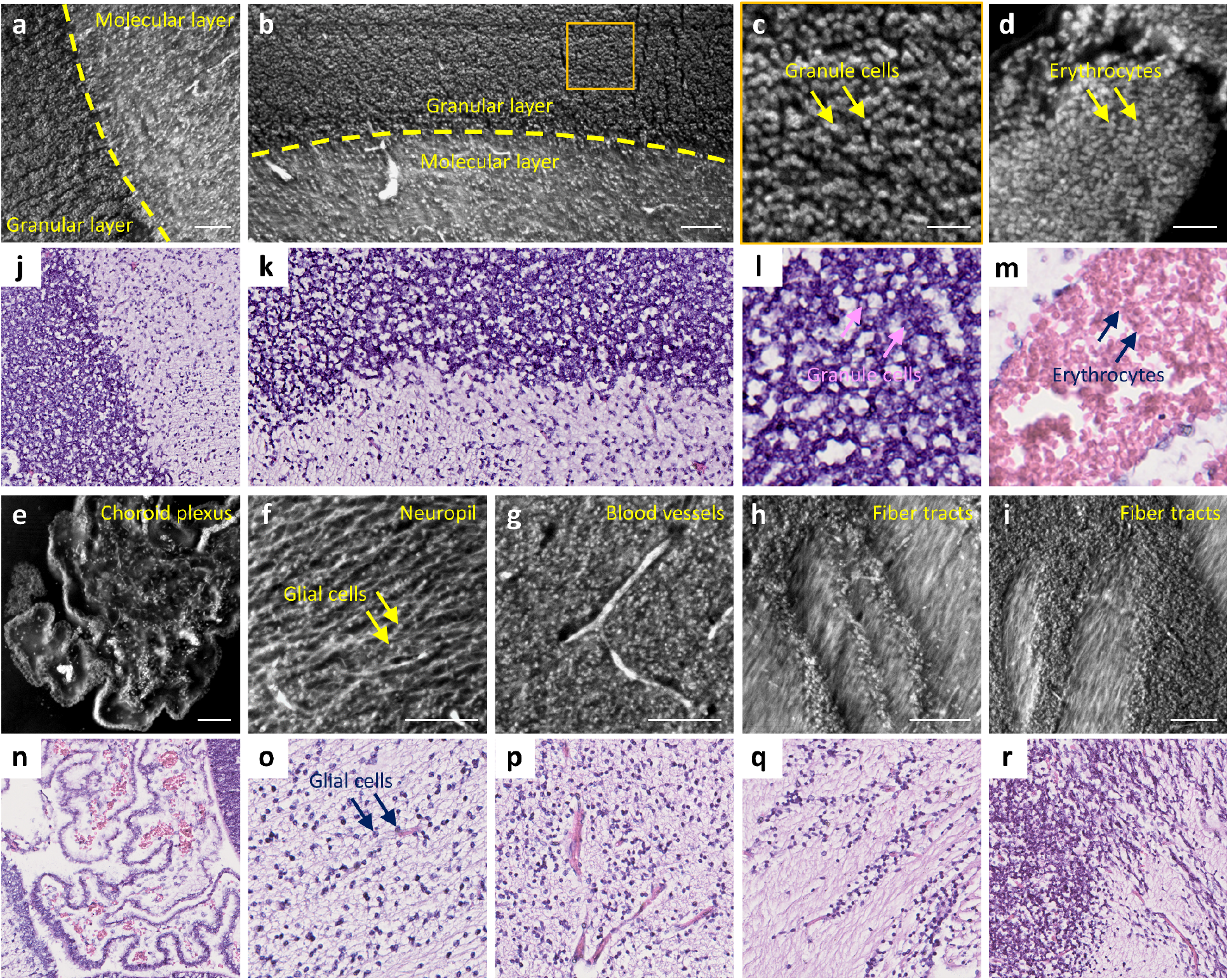
Image gallery of features extracted from label-free MATE images of the human brain block. **a,b**, Cerebellar cortex, with the boundary between the granular layer and molecular layer marked by yellow dashed lines. **c**, A zoomed-in MATE image of the orange solid region in b. **d**, Erythrocytes. **e**, Choroid plexus. **f**, Neuropil. **g**, Blood vessels. **h,i**, Fiber tracts. **j–r**, The corresponding features extracted from the H&E-stained histological images. All features are shown in a coronal view. Scale bars, 30 μm (c,d) and 100 μm (a,b,e–i).

### MATE imaging of an FFPE human gallbladder block

Finally, an FFPE human gallbladder block is imaged by MATE (Fig. 6a–d). Similarly, the block is consecutively sectioned and stained to obtain the corresponding H&E-stained images for comparison (Fig. 6e–h). Despite that tissue deformation occurs during microtome sectioning and slide mounting, molecular features such as erythrocytes (Fig. 6b,f) in the veins (Fig. 6c,g), and anatomical features such as perimuscular connective tissue (Fig. 6d,h) are well characterized in both MATE and H&E-stained images with high fidelity, validating the feasibility of using MATE to image different organs.

**Figure 6.**
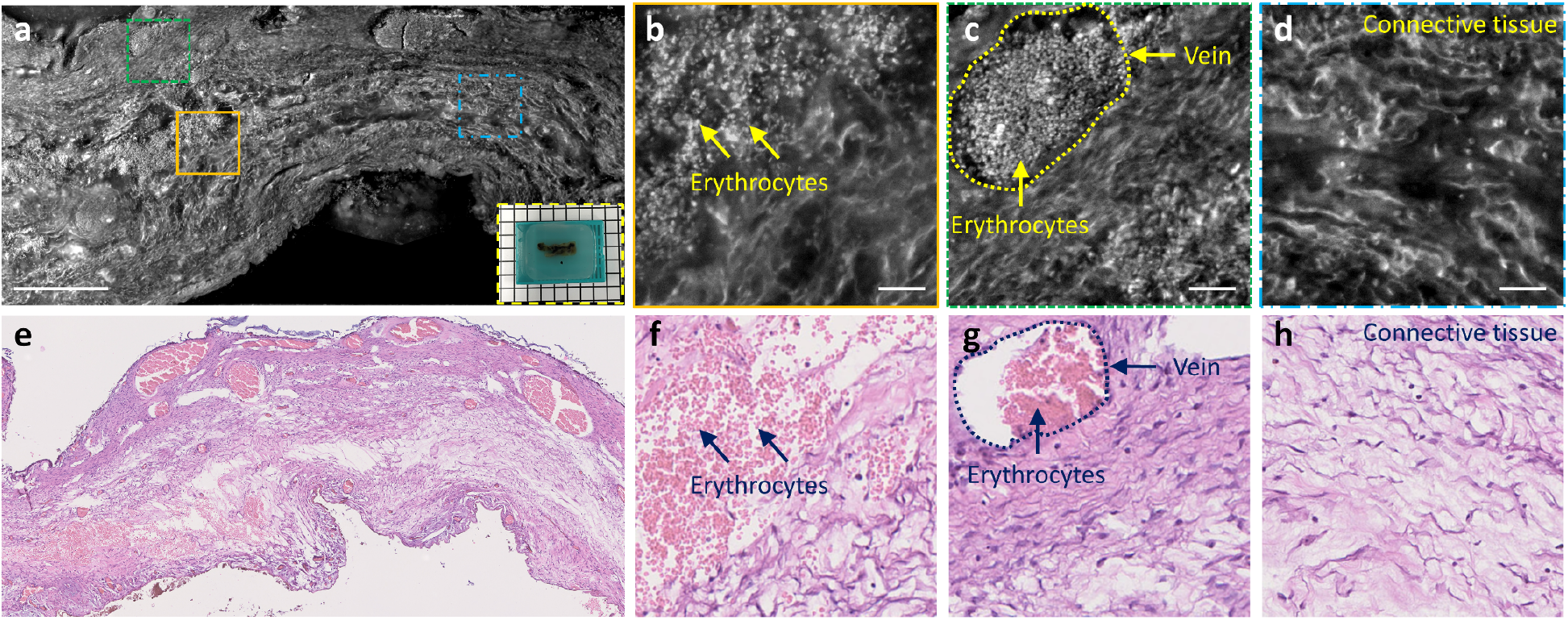
MATE imaging of an FFPE human gallbladder block. **a**, A MATE image of an FFPE human gallbladder block, inset at the bottom right shows the photograph of the specimen. **b–d**, Zoomed-in MATE images of orange solid, green dashed, and blue dashed regions in a, respectively. **e–h**, The corresponding H&E-stained histological images. Scale bars, 500 μm (a) and 50 μm (b–d).

## Discussion

MATE is a promising and transformative 3D histological imaging platform which enables rapid and comprehensive histopathological analysis of complex whole organs without tissue staining or clearing. However, there are still deviations between MATE and H&E-stained histological images. First, the nucleoli structures are better observed in H&E-stained images than that in MATE under the same magnification (e.g., Supplementary Figs. 2i,l and 3b,h). This is likely because the fluorescence property of nucleoli in the detected spectral range is not chemically identical to the histological stains. Second, the texture of fiber tracts in the white matter of the human brain is better visualized in MATE than that in H&E-stained images (e.g., Fig. 5h,i,q,r). This is possibly due to the proteins in these fibrous structures present a high quantum yield under deep-UV excitation while eosin exhibits similar affinity across the cytoplasm.

Intrinsic fluorescence excited by deep-UV light naturally forms a contrast mechanism for high-content label-free imaging, but it varies significantly among different types of tissues/biomolecules. In addition, fluorescence properties of endogenous fluorophores, such as excitation/emission spectra and quantum yields, are highly related to the biochemical environment and metabolic status. Therefore, some brightness variations can be observed in MATE images. For example, the cell nuclei show a negative contrast in mouse brain blocks (Fig. 3 and Supplementary Fig. 3) while features a positive contrast in the human brain block (Fig. 5). Besides, the fluorescence from erythrocytes, generated by photodecomposition of hemoglobin with deep-UV irradiation, is sensitive to reactive oxygen species (37), thus presenting different fluorescence intensity among thin tissue slices (denoted by asterisks in Supplementary Fig. 2). MATE allows simultaneous visualization of cell nuclei, fiber tracts, and blood vessels by non-specific excitation of various biomolecules, posing a challenge to accurate segmentation of clinically relevant image features. We believe that this problem can be solved by using advanced neural networks, which would enable a more faithful tissue analysis (38).

MATE is currently in an early stage of development. The technical improvements can be further realized in terms of imaging speed and spatial resolution. MATE enables 3D reconstruction of 1-cm^3^ tissue volume at 1.2 × 1.2 × 10 μm^3^ voxel resolution within 4 hours in the current setting. The acquisition process can be further accelerated by (1) employing multiple high-power UV-LEDs to minimize the camera integration time for each image tile, (2) using high-speed motorized stages for fast mechanical sectioning, and (3) applying contour detection to avoid unnecessary background acquisitions. In addition, UV absorption from the embedded tissues is the current limiting factor for axial resolution, which is measured to be ~10 μm in our experiments. By integrating with structured illumination microscopy(21), or doping the sample embedding media with strong UV-absorbing dyes(32), the axial resolution is expected to be enhanced by an order of magnitude, facilitating diverse applications where require high voxel resolution such as long-range axon tracking and capillary network mapping in whole-brain imaging. Furthermore, virtually-stained MATE images can be generated with the assistance of unsupervised learning (39, 40), eliminating any training for pathologists in image interpretation for diagnostic decision-making.

In summary, we developed a label-free, automated, and ready-to-use 3D imaging technique that can be used routinely for comprehensive 3D histopathological analysis of complex and bulky tissues. With the combination of block-face imaging and serial microtome sectioning, MATE enables rapid and label-free imaging of paraffin-embedded whole organs at an acquisition speed of 1 cm^3^ per 4 hours with a voxel resolution of 1.2 × 1.2 × 10 μm^3^. We showed that different anatomical structures, including cell nuclei, fiber tracts, and blood vessels in mouse/human brains, can be simultaneously visualized in MATE without tissue staining or clearing. In addition, diagnostic features, such as nuclear size and packing density, can be quantitatively extracted from MATE with high accuracy. As an augment to the current slide-based 2D histology, MATE holds great promise for studying a variety of disease processes in 3D, which should be carried out as follow-up work.

## Method

### Collection of biological tissues

C57BL/6 mice were euthanized by CO_2_. The brain tissue was then harvested from the mice immediately and fixed in 4% neutral-buffered formalin at room temperature for 24 hours, followed by standard FFPE tissue preparation protocol and paraffin-embedded as block specimens for MATE imaging. To prepare thin tissue slices for validation, the block specimens were sectioned at the surface by a microtome (RM2235, Leica Microsystems Inc.) with 7-μm thickness and stained by H&E, and subsequently imaged by a digital slide scanner (NanoZoomer-SQ, Hamamatsu Photonics K.K.) to generate the corresponding histological images. All experiments were carried out in conformity with a laboratory animal protocol approved by the Health, Safety and Environment Office of Hong Kong University of Science and Technology (HKUST). The human sample protocol was approved by the Institutional Review Board at the Prince of Wales Hospital. The imaged tissue was considered as leftover tissue, i.e., it represents a portion of a collected specimen that is not needed for assessment of diagnostic, prognostic, and other parameters in the diagnosis and treatment of the patient.

### Configuration of the MATE system

As shown in Fig. 1, the MATE system is configured in a reflection mode to accommodate tissues with any size and thickness. A UV-LED (285-nm wavelength, M285L5, Thorlabs Inc.) is spectrally filtered by a band-pass filter (FF01-285/14-25, Semrock Inc.) and obliquely focused onto the bottom surface of the specimen through a pair of relay lenses (#33-957, Edmund Optics Inc., and LA4148-UV, Thorlabs Inc.). The excitation power is measured to be 100 mW with an illumination area of ~1 mm^2^. Oblique illumination circumvents the use of any UV-transmitting optics and fluorescence filters because the backscattered UV light is naturally blocked by the glass objective and tube lens which are spectrally opaque at 285 nm. The UV-excited intrinsic fluorescence is detected by an inverted microscope which consists of a plan achromat objective lens (Plan Fluor, 10×/0.3 NA, Olympus Corp.) and an infinity-corrected tube lens (TTL180-A, Thorlabs Inc.), and subsequently imaged by a monochrome scientific complementary metal-oxide-semiconductor camera (PCO edge 4.2, 2048 × 2048 pixels, 6.5-μm pixel pitch, PCO. Inc.) which can theoretically reach an imaging throughput of ~800 megabytes/s. The paraffin-embedded specimen is clamped towards the objective lens by an adjustable sample holder, and rigidly mounted on a 3-axis motorized stage (L-509.20SD00, 1-μm bidirectional repeatability, PI miCos GmbH) to control both *x-y* scanning at 1.2-mm interval for in-plane mosaic imaging, and *z* scanning at the 10-μm interval for microtome sectioning. The lab-built microtome is tuned with a cutting angle of ~30°, and placed conjugated with the objective’s focal plane to obtain sharp in-focus images. The thin tissue slices are sliced off and dropped down due to the gravity and collected in a collection area. To balance acquisition speed, image quality, and sectioning stability, the motorized stage is translated at a velocity of 15 mm/s for microtome sectioning, and the acquisition time for each in-plane image tile is set to 110 ms (100-ms camera integration time plus 10-ms stage settling time). The captured image tiles are stitched during image acquisition, which is synchronized with motor scanning via our lab-designed LabVIEW software (National Instruments Corp.) and triggering circuits. The system is fully automated to provide a series of distortion- and registration-free images with intrinsic absorption-based contrast, enabling rapid and label-free 3D imaging of paraffin-embedded whole organs at an acquisition speed of 1 cm^3^ per 4 hours. The lateral resolution of MATE was characterized by imaging 200-nm-diameter fluorescent beads (B200, emission at 445 nm, Thermo Fisher Scientific Inc.), and the full width at half maximum is measured to be 1.2 μm (Supplementary Fig. 1).

### Image analysis

MATE and H&E-stained histological images were segmented by a free Fiji plugin (trainable Weka segmentation), and subsequently binarized and analyzed in Fiji to acquire the cross-sectional area and centroid of each cell nucleus. With the localized center positions of cell nuclei, the intercellular distance was calculated to be the shortest adjacent distance to a neighboring cell nucleus. To generate a nuclear density map, the MATE images were processed by a Jerman’s spherical enhancement filter(33), and subsequently binarized and analyzed in Fiji to localize each cell nucleus. After that, the center positions of the cell nuclei in the image were set to unit-amplitude against a zero-amplitude background, in which cell counting was performed in every 50 μm × 50 μm surrounding area and normalized to generate the nuclear density map (detailed method shown in Supplementary Fig. 5). The serial registration-free block-face images were processed and rendered in Avizo software (Thermo Fisher Scientific Inc.) for 3D visualization.

## Data availability

All data involved in this work, including raw/processed images provided in the manuscript and Supplementary Information, are available from the corresponding author upon request.

## Acknowledgments

The Translational and Advanced Bioimaging Laboratory (TAB-Lab) at HKUST acknowledges the pathology department from the Prince of Wales Hospital for providing the human samples. We also acknowledge the support of the Hong Kong Innovation and Technology Commission (ITS/036/19) and the Research Grants Council of the Hong Kong Special Administrative Region (26203619 and 16208620).

## Author contributions

Y.Z., L.K., and T.T.W.W. conceived of the study. Y.Z. and L.K. built the imaging system. Y.Z., L.K., and V.T.C.T prepared the specimens involved in this study. Y.Z. performed imaging experiments. L.K. performed histological staining. Y.Z. processed and analyzed the data. Y.Z. and T.T.W.W. wrote the manuscript. T.T.W.W. supervised the whole study.

