## Supplementary Information for "Three-dimensional label-free histological imaging of whole organs by microtomy-assisted autofluorescence tomography"

The Supplementary Material includes:

Supplementary Figure 1 | Experimental characterization of MATE's lateral resolution.

Supplementary Figure 2 | MATE imaging of FFPE thin tissue slices.

Supplementary Figure 3 | MATE imaging of an FFPE mouse brain block.

Supplementary Figure 4 | Imaging depth of MATE in an FFPE mouse brain block.

Supplementary Figure 5 | Calculation of nuclear density map.

Supplementary Figure 6 | Effect of different embedding materials.

Other Supplementary Material for this manuscript includes the following:

Supplementary Video 1 | Close-up of MATE and H&E-stained images of an FFPE mouse brain block.

Supplementary Video 2 | A series of coronal sections of an intact FFPE mouse brain imaged by MATE.

Supplementary Video 3 | A series of coronal sections of an FFPE human brain block imaged by MATE.

Supplementary Video 4 | High-resolution 3D model of fiber pathways in the white matter of the human brain.

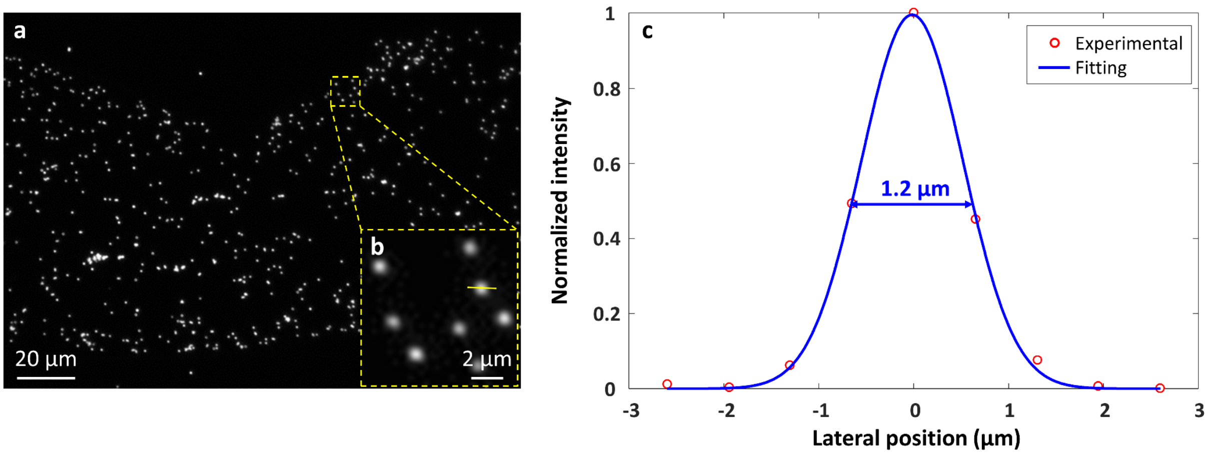


**Supplementary Figure 1 | Experimental characterization of MATE's lateral resolution. a**, A MATE image of blue fluorescent beads (200-nm in diameter with an emission wavelength of 445 nm). **b**, A zoomed-in MATE image of the yellow dashed box in a. **c**, Gaussian-fitted intensity distribution along the solid line in b, showing that the lateral resolution is 1.2 µm.

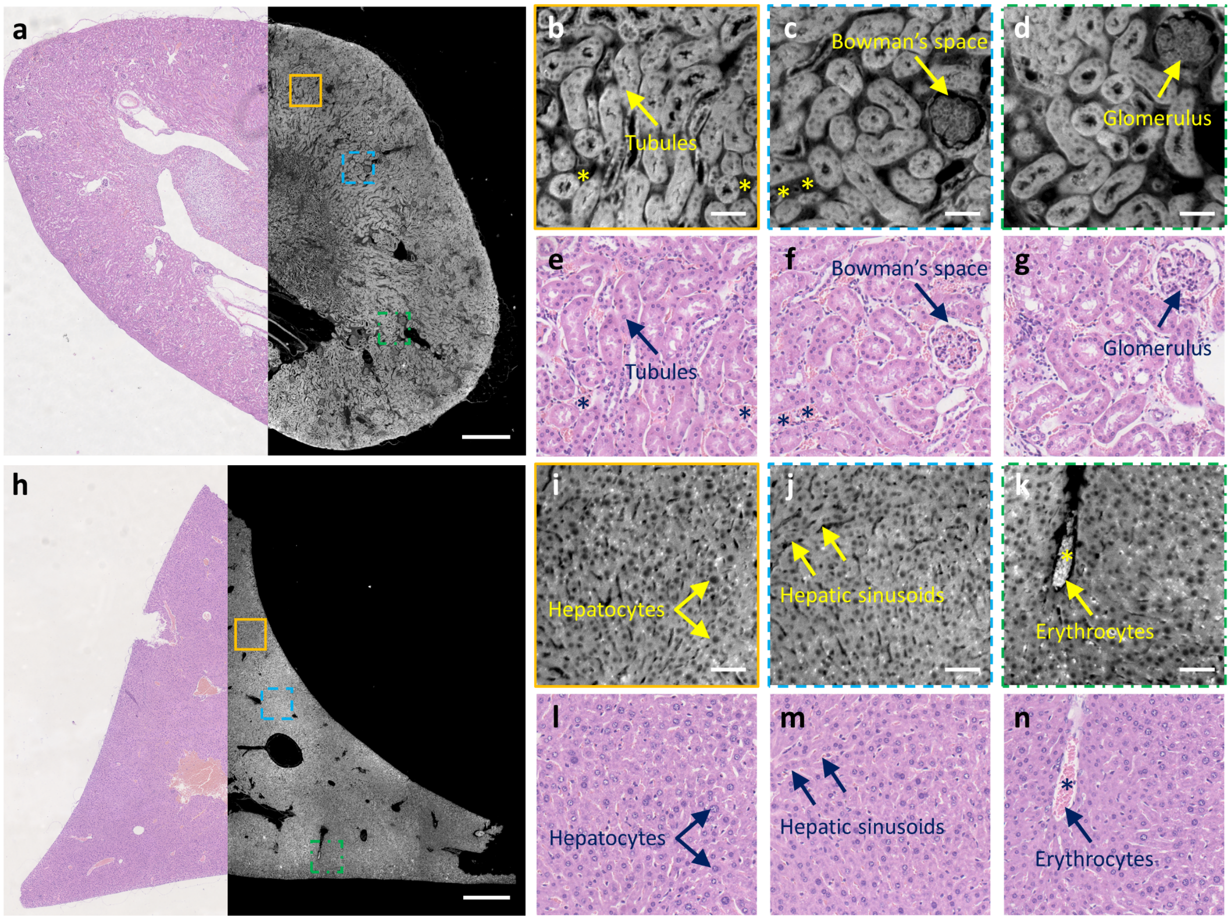


**Supplementary Figure 2 | MATE imaging of FFPE thin tissue slices. a,** A combined MATE and H&E-stained mosaic image of a mouse kidney section. **b**–**d,** Zoomed-in MATE images of orange solid, blue dashed, and green dashed regions in a, respectively. **e**–**g,** The corresponding H&E-stained images. **h,** A combined MATE and H&E-stained mosaic image of a mouse liver section. **i**–**k,** Zoomed-in MATE images of orange solid, blue dashed, and green dashed regions in h, respectively. **l**–**n,** The corresponding H&E-stained images. Intensity variations of erythrocytes are denoted by the asterisks. Scale bars, 500 µm (a,h) and 50 µm (b–d, i–k).

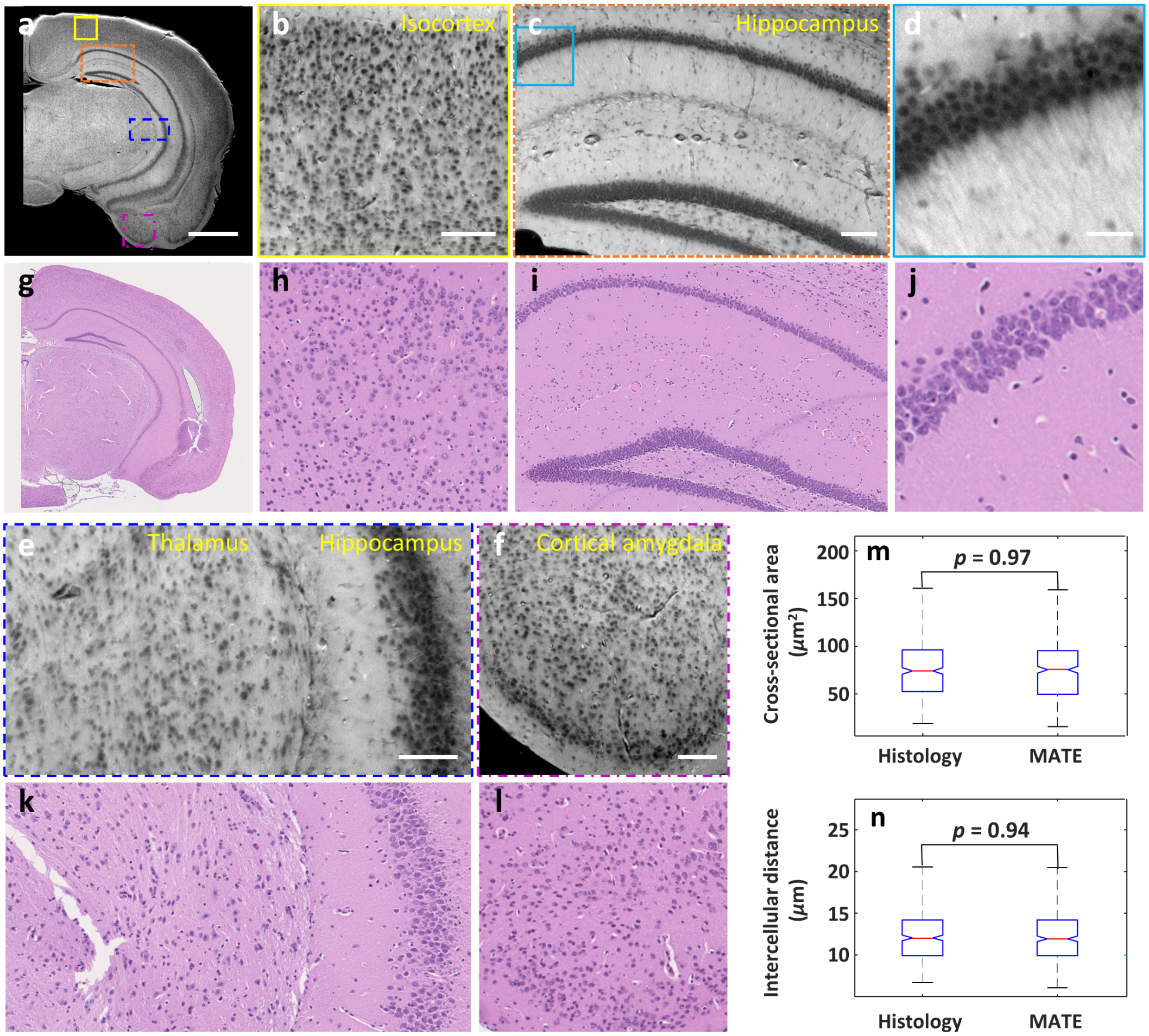


**Supplementary Figure 3 | MATE imaging of an FFPE mouse brain block. a,** A MATE image of an FFPE mouse brain block. **b,c,** Zoomed-in MATE images of yellow solid and orange dashed regions in a, respectively. **d,** A zoomed-in MATE image of the blue solid region in c. **e,f,** Zoomed-in MATE images of blue dashed and magenta dashed regions in a, respectively. **g–l,** The corresponding H&E-stained images. **m,n,** Distributions of cross-sectional area and intercellular distance extracted from b and h. Scale bars, 1 mm (a), 100 μm (b,c,e,f), and 30 μm (d).

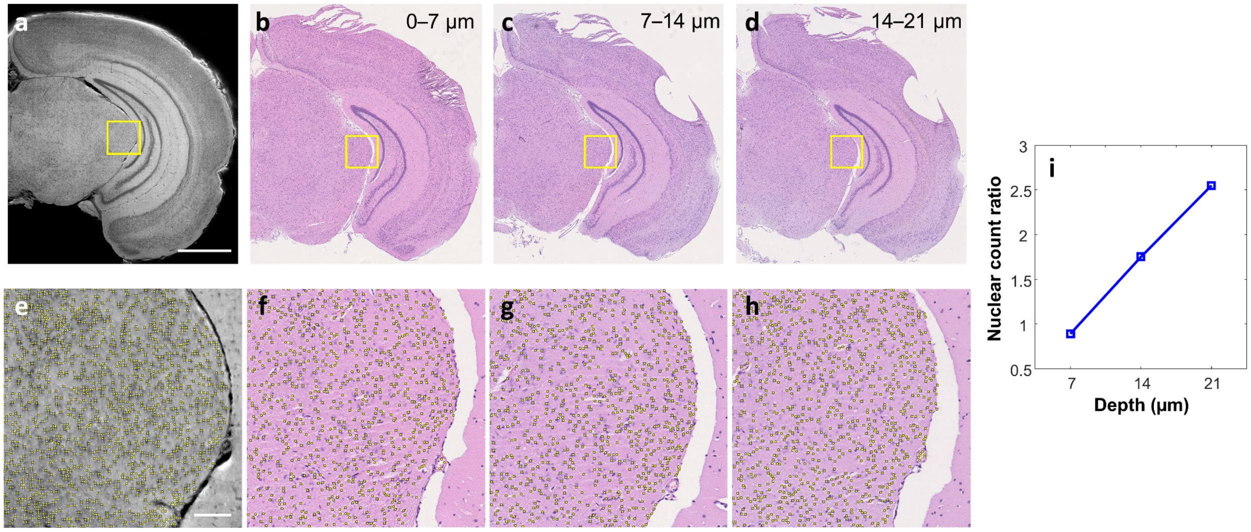


**Supplementary Figure 4 | Imaging depth of MATE in an FFPE mouse brain block. a,** A MATE image of an FFPE mouse brain block. **b**–**d,** H&E-stained images of thin mouse brain slices which are consecutively sectioned from the block surface with 7-μm thickness. **e**–**h,** Zoomed-in images of yellow solid regions in a–d, respectively. **i,** The ratio of the nuclear count in the H&E-stained images within a given depth range to that in the MATE image. Scale bars, 1 mm (a) and 100 μm (e).

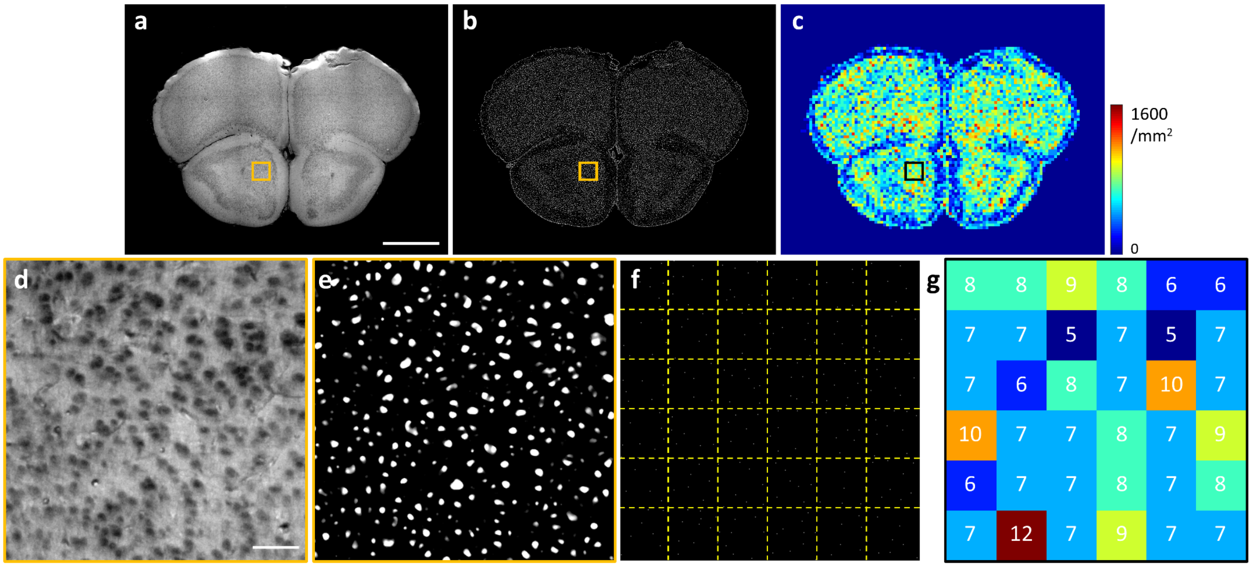


**Supplementary Figure 5 | Calculation of nuclear density map. a,** A representative MATE image from the mouse brain dataset. **b,** A masked MATE image processed by a Jerman's spherical enhancement filter. **c,** Nuclear density map calculated from b. **d,e,** Zoomed-in images of the marked regions in a and b, respectively. **f,** A binary image that contains center positions of all nuclei in d. **g,** A zoomed-in image of the marked region in c, with the cell counting performed in every 50 µm × 50 µm surrounding area indicated by yellow dashed grids in f. Scale bars, 1 mm (a) and 50 µm (d).

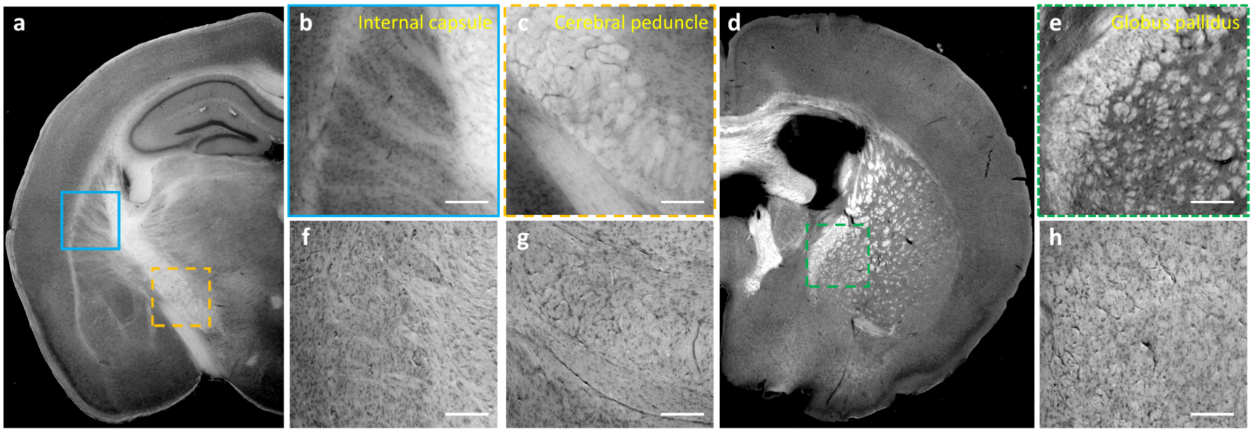


**Supplementary Figure 6 | Effect of different embedding materials. a,** A MATE image of an agarose-embedded mouse brain tissue with 200-μm thickness. **b,c,** Zoomed-in MATE images of the blue solid and orange dashed regions in a, respectively. **d,** Another MATE image of an agarose-embedded mouse brain tissue with 200-μm thickness. **e,** A zoomed-in MATE image of the green dashed region in d. **f**–**h**, The corresponding features extracted from a paraffin-embedded mouse brain block. Scale bars, 200 μm.
